# A cortical hierarchy of localized and distributed processes revealed via dissociation of task activations, connectivity changes, and intrinsic timescales

**DOI:** 10.1101/262626

**Authors:** Takuya Ito, Luke J. Hearne, Michael W. Cole

## Abstract

Many studies have identified the role of localized and distributed cognitive functionality by mapping either local task-related activity or distributed functional connectivity (FC). However, few studies have directly explored the relationship between a brain region’s localized task activity and its distributed task FC. Here we systematically evaluated the differential contributions of task-related activity and FC changes to identify a relationship between localized and distributed processes across the cortical hierarchy. We found that across multiple tasks, the magnitude of regional task-evoked activity was high in unimodal areas, but low in transmodal areas. In contrast, we found that task-state FC was significantly reduced in unimodal areas relative to transmodal areas. This revealed a strong negative relationship between localized task activity and distributed FC across cortical regions that was associated with the previously reported principal gradient of macroscale organization. Moreover, this dissociation corresponded to hierarchical cortical differences in the intrinsic timescale estimated from resting-state fMRI and region myelin content estimated from structural MRI. Together, our results contribute to a growing literature illustrating the differential contributions of a hierarchical cortical gradient representing localized and distributed cognitive processes.

**Highlights:**

- Task activations and functional connectivity changes are negatively correlated across cortex
- Task activation and connectivity dissociations reflect differences in localized and distributed processes in cortex
- Differences in localized and distributed processes are associated with differences in intrinsic timescale organization
- Differences in localized and distributed processes are associated with differences in cortical myelin content
- Cortical heterogeneity in localized and distributed processes revealed by activity flow mapping prediction error

## Introduction

The brain processes information in both a localized and distributed manner. At the macroscale, localized functionality has typically been revealed by measuring the local activity of brain regions in response to experimental conditions (Poldrack, 2011; Wallis, 2018). In contrast, distributed neural processing across large-scale cortical systems has typically been studied by measuring the task-state functional connectivity (FC; i.e., covariation of neural signal across different brain regions) (Amico et al., 2019; Cole et al., 2014, 2013; Gratton et al., 2016; Hermundstad et al., 2013; Ito et al., 2019b; Krienen et al., 2014). Yet few studies have identified how these two processes – localized and distributed – differ across the cortical hierarchy during cognition. Here we directly assess the differential roles of distributed and localized processing by characterizing regional differences in local task activations and distributed task FC during multiple task states. Further, we tie these regional differences to two intrinsic properties of hierarchical cortical organization: intrinsic timescale organization estimated from resting-state functional magnetic resonance imaging (fMRI) and regional myelin content from structural MRI.

Many studies have successfully ascribed cognitive functions onto specific brain regions, demonstrating localized cognitive processes (Genon et al., 2018; Norman et al., 2006; Poldrack, 2011). Such studies focus on capturing task-/stimulus-evoked neural response patterns related to the cognitive processes recruited by experimental paradigms. This endeavor has identified sets of brain regions associated with, for example, visual (Haxby et al., 2006; Hubel and Wiesel, 1962; Kanwisher et al., 1997), language (Fedorenko et al., 2012; Huth et al., 2016), and motor processes (Churchland et al., 2006; Yokoi and Diedrichsen, 2018), indicating a wealth of functional diversity across the cortex.

Research focused on the distributed nature of cognitive processing typically characterize large-scale functional network organization during task manipulation (Amico et al., 2019; Chauvin et al., 2019; Cole et al., 2014, 2013; Gratton et al., 2016; Ito et al., 2019a; Krienen et al., 2014; Shine et al., 2016). Such studies have shown that though large-scale functional network changes are often task-dependent, transmodal areas may be disproportionately involved in distributed processes relative to unimodal areas (Cole et al., 2013; Gratton et al., 2016; Hwang et al., 2018). While task-evoked activations have been shown to be negatively associated with FC changes across brain regions (Ito et al., 2019a), it is unclear whether this local versus distributed dichotomy is related to intrinsic macroscale cortical organization.

Recent studies from our group illustrated that regional task-evoked activity can be predicted from a distributed activity flow process (Cole et al., 2016; Ito et al., 2017). This approach – activity flow mapping – assumes that task-evoked activity is propagated between brain regions through distributed functional connections that can be estimated during resting-state fMRI. While the success of this approach assumes that local activity is the result of a distributed neural process, there are likely gradients of functional heterogeneity in the brain (Demirtaş et al., 2019; Huntenburg et al., 2018; Yeo et al., 2015), suggesting that cortical regions may not behave in a uniformly distributed manner. Previous work has shown that that there are structural and functional bases of hierarchical cortical heterogeneity, such as in regional myelin content (which has been used as a proxy to index anatomical hierarchy) and intrinsic timescale organization (Burt et al., 2018; Felleman and Van Essen, 1991; Murray et al., 2014; Wang, 2020). Murray et al. illustrated that lower-order cortical areas tend to operate at fast timescales, potentially supporting stimulus-/task-locked activity, while higher-order cortical areas tend to operate at slower timescales, potentially supporting information integration from lower-order areas (Murray et al., 2014). However, those conclusions were based on non-human primate electrophysiology data. Considering this, we sought to characterize hierarchical timescale organization in human fMRI data and evaluate whether it was related to local and distributed processes during cognitive task processing.

We report empirical evidence demonstrating a dissociation of local and distributed processes across the cortical hierarchy during task states. We extended previous results to demonstrate that the dissociation of task-evoked local activity and distributed FC (Ito et al., 2019a) is related to hierarchical cortical organization. Specifically, we first report cortical heterogeneity in the intrinsic timescales estimated during resting-state fMRI (Lurie and D’Esposito, 2019; Murray et al., 2014), finding that regions with faster intrinsic timescales had strong activations and reduced FC during task states. Second, we show that regional myelin content (T1w/T2w scans), which has recently been shown to be associated with local genomic, structural, and biophysical properties (Burt et al., 2018; Demirtaş et al., 2019), also explains cortical differences in local task activations and distributed FC during multiple task states. Finally, we show that higher-order transmodal regions are better predicted via activity flow mapping relative to unimodal regions (Cole et al., 2016), indicating that the activity of these regions reflect more distributed processes rather than local processes. Together, our results illustrate the differential contributions of localized and distributed cognitive processing along a hierarchical cortical gradient.

## Materials and Methods

### Data and paradigm

We use a publicly available Human Connectome Project (HCP) data set (Van Essen et al., 2013). The details below are identical to those reported in (Ito et al., 2019a), and are included below.

The Rutgers University institutional review board approved this study. We obtained data from the Washington University-Minnesota Consortium of the HCP [31]. We used 352 subjects from the HCP 1200 release for empirical analyses. Details and procedures of subject recruitment can be found in (Van Essen et al., 2013). The 352 subjects were selected based on: quality control assessments (i.e., any participants with any quality control flags were excluded, including 1) focal anatomical anomaly found in T1w and/or T2w scans, 2) focal segmentation or surface errors, as output from the HCP structural pipeline, 3) data collected during periods of known problems with the head coil, 4) data in which some of the FIX-ICA components were manually reclassified; low-motion participants (i.e., exclusion of participants that had any fMRI run in which more than 50% of TRs had greater than 0.25mm framewise displacement); removal according to family relations (unrelated participants were selected only, and we excluded those with no genotype testing).

All participants were recruited from Washington University in St. Louis and the surrounding area. The 352 subjects were split into two equal cohorts of 176 subjects (99 females and 84 females). The exploratory cohort had a mean age of 29 years of age (range=22-36 years of age), and the replication cohort had a mean age of 28 years of age (range=22-36 years of age). All subjects gave signed, informed consent in accordance with the protocol approved by the Washington University institutional review board. Whole-brain multiband echo-planar imaging acquisitions were collected on a 32-channel head coil on a modified 3T Siemens Skyra with TR=720 ms, TE=33.1 ms, flip angle=52°, Bandwidth=2,290 Hz/Px, in-plane FOV=208×180 mm, 72 slices, 2.0 mm isotropic voxels, with a multiband acceleration factor of 8. The HCP collected data over two days for each subject. On the first day, anatomical scans were collected (including T1-weighted and T2-weighted images acquired at 0.7 mm isotropic voxels) followed by two resting-state fMRI scans (each lasting 14.4 minutes), and ending with a task fMRI component. The second day consisted of collecting a diffusion imaging scan, followed another set of two resting-state fMRI scans (each lasting 14.4 minutes) and a task fMRI session.

Each of the seven tasks was collected over two consecutive fMRI runs. The seven tasks consisted of an emotion cognition task, a gambling reward task, a language task, a motor task, a relational reasoning task, a social cognition task, and a working memory task. Briefly, the emotion cognition task required making valence judgements on negative (fearful and angry) and neutral faces. The gambling reward task consisted of a card guessing game, where subjects were asked to guess the number on the card to win or lose money. The language processing task consisted of interleaving a language condition, which involved answering questions related to a story presented aurally, and a math condition, which involved basic arithmetic questions presented aurally. The motor task involved asking subjects to either tap their left/right fingers, squeeze their left/right toes, or move their tongue. The reasoning task involved asking subjects to determine whether two sets of objects differed from each other in the same dimension (e.g., shape or texture). The social cognition task was a theory of mind task, where objects (squares, circles, triangles) interacted with each other in a video clip, and subjects were subsequently asked whether the objects interacted in a social manner. Lastly, the working memory task was a variant of the N-back task.

Further details on the resting-state fMRI portion can be found in (Smith et al., 2013), and additional details on the task fMRI components can be found in (Barch et al., 2013).

### Preprocessing

Preprocessing details below follow identical procedures from (Ito et al., 2019a), and are described below.

Minimally preprocessed data were obtained from the publicly available HCP data. Minimally preprocessed surface data was downsampled into 360 parcels using the (Glasser et al., 2016) atlas. We additionally preprocessed on the parcellated data for resting-state fMRI and task state fMRI. This included removing the first five frames of each run, de-meaning and de-trending the time series, and performing nuisance regression on the minimally preprocessed data (Ciric et al., 2017). Nuisance regression removed motion parameters and physiological noise. Specifically, six primary motion parameters were removed, along with their derivatives, and the quadratics of all regressors (24 motion regressors in total). We applied aCompCor on the physiological time series extracted from the white matter and ventricle voxels (5 components each) (Behzadi et al., 2007). In addition, we included the derivatives of each component, and the quadratics of all physiological noise regressors (40 physiological noise regressors in total). The combination of motion and physiological noise regressors totaled 64 nuisance parameters and is a variant of previously benchmarked nuisance regression models reported in (Ciric et al., 2017).

We did not apply global signal regression (GSR), given that GSR artificially induces negative correlations (Murphy et al., 2009). We included aCompCor as a preprocessing step given that aCompCor does not include GSR while including some of its benefits (some extracted components are highly similar to the global signal) (Power et al., 2018). This approach is conceptually similar to temporal-ICA-based artifact removal procedure that seeks to remove global artifact without removing global neural signals, which contains behaviorally relevant information such as vigilance (Glasser et al., 2018; Wong et al., 2013). Further, we included the derivatives and quadratics of each component time series (within aCompCor) to further reduce artifacts. Code to perform this regression is publicly available online using python code (version 2.7.15) (https://github.com/ito-takuya/fmriNuisanceRegression).

Data for task FC analyses were additionally preprocessed using a standard general linear model (GLM). We fitted the task timing (block design) for each task condition using a finite impulse response (FIR) model (with a lag extending to 25 TRs after task block offset) to remove the mean evoked task-related activity (Cole et al., 2019). (Across seven tasks, there were 24 task conditions.) Removing the mean task-evoked response (i.e., main effect of task) is critical to isolate the spontaneous neural activity, and has been performed in the spike count correlation literature for decades (Aertsen et al., 1989; Cohen and Kohn, 2011).

### Task activation analysis

We performed a standard task GLM analysis on fMRI task data to evaluate the task-evoked activity. The task timing for each of the 24 task conditions was convolved with the SPM canonical hemodynamic response function to obtain task-evoked activity estimates (Friston et al., 1994). FIR modeling was not used when modeling task-evoked activity. Coefficients were obtained for each parcel in the Glasser et al. (2016) cortical atlas for each of the 24 task conditions.

To characterize the degree of local processing within a region, we characterized the average activation magnitude across multiple tasks. The calculation was identical to a previous report (He, 2013). Specifically, for each condition, we obtained the absolute magnitude of the t-statistic relative to 0 across subjects. (This characterized the magnitude of the task-evoked activity relative to baseline, independent of sign.) This was then averaged across task conditions, resulting in a 360×1 vector of task activation magnitudes across multiple task conditions. This vector was used to characterize the average degree of local processing across multiple tasks.

### Functional connectivity analyses

We computed the task state FC across all task conditions (between all pairs of brain regions) after removing the mean task-evoked response for each condition separately. This resulted in a single 360×360 FC matrix. To obtain a statistically comparable resting-state FC matrix (with equivalent temporal intervals), we applied the identical calculation to resting-state data. This involved first regressing out the same task design matrix used during task-state regression in resting-state data. This was possible given that the number of timepoints of the combined resting-state scans in the HCP data set exceeded the number of timepoints of the combined task-state scans (4800 resting-state TRs > 3880 task-state TRs). We then obtained each region’s resting-state FC matrix by applying the same task block design onto resting-state data (i.e., ensuring correlations were obtained using the same temporal intervals as task data, though data was from resting state).

To obtain the average FC strength change from rest to task for each brain region, we subtracted weighted degree centrality (also often referred to as global brain connectivity) computed from task data from weighted degree centrality computed from rest data (Cole et al., 2010; Rubinov and Sporns, 2010). This resulted in a 360×1 vector of the averaged FC strength change from rest to task across multiple tasks.

### Peak activation approach for calculating task activations and FC changes

We conducted an additional set of analyses that did not involve removing the mean task-evoked response (via task regression) to ensure results were not dependent on this step. In principle the mean task-evoked response removal step is critical for avoiding false positives (Cole et al., 2019), yet we identified an alternate way to avoid these false positives: focusing on event-to-event/block-to-block variance and covariance. This involved taking the peak absolute value of a time series during each task block (after baselining the time series to the mean activity level across all inter-block rest periods). The extracted data points reflected the peak task-evoked activation within each block for each region. The peak activation values were then summarized by averaging across blocks for task activation level estimation, and task FC was computed as correlations across these block-to-block peak activations between regions. This is conceptually similar to estimating the beta coefficient for each task block separately (Rissman et al., 2004), but we instead estimate the peak of the block (rather than the coefficient for the entire block).

To estimate an equivalent resting-state FC that controlled for the number of time point samples (i.e., samples across ‘blocks’), we applied the same procedure to resting-state data. Specifically, we applied the same task design matrix onto resting-state data to identify temporally equivalent ‘pseudo-blocks’, and extracted the peak absolute BOLD value of each pseudo-block period. Resting-state FC was then computed as the correlation across blocks. Task-state FC change for each region was calculated as the difference between task and rest FC values.

### Intrinsic timescale

We estimated the intrinsic timescales of brain regions using resting-state fMRI. We computed the autocorrelation function of each brain region. The main results are reported by estimating the autocorrelation function with a lag of 100 timepoints (72s). Results were reproduced using shorter lags (e.g., 40 and 50 time points). To estimate the intrinsic timescale, we fitted a nonlinear exponential decay function with an offset to the empirically estimated autocorrelation function as previously described in (Murray et al., 2014). The exponential decay function was fit as a function of the time lag *k*Δ between time bins *k* = |*i* − *j*|, and obeyed the equation

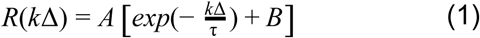

where *A* corresponds to a scaling factor, *B* reflects the offset for contribution of timescales longer than the observation window, and τ corresponds to the intrinsic timescale (i.e., rate of decay). This procedure was performed for every parcel separately using a nonlinear least-squares fitting procedure using the ‘Trust Region Reflective’ algorithm as implemented in scipy.optimize.curvefit (version 1.2.1; python version 3.7.3). Given the biological implausibility of a negative scaling factor (*A*) and negative intrinsic timescale (τ), we constrained our solution using parameter bounds *A* ∈ [0, ∞) and *B* ∈ (− ∞, ∞) and τ ∈ [0, ∞).

### Cortical myelin map

Cortical myelin maps were obtained in surface-based CIFTI file format from the authors of a previously published study (Burt et al., 2018), and are briefly described below. T1w/T2w maps were obtained from the (HCP) (Van Essen et al., 2013). T1w and T2w maps were registered to a standard reference space (MNI) using an areal-feature-based technique (Glasser et al., 2016; Robinson et al., 2014). Cortical T1w/T2w maps were averaged across 339 subjects (see Burt et al. 2018 for additional details).

Parcellated maps were obtained by downsampling the surface-based CIFTI file into 360 cortical regions using the Glasser et al. (2016) atlas. Previous work has shown that this T1w/T2w contrast reflects cortical myelin content (Glasser and Van Essen, 2011), and that the parcellated maps are highly stable across individual subjects. (The mean pairwise Spearman rank correlation between subjects’ individual maps was previously found to be rho=0.94 (see Burt et al., 2018).)

### Activity flow mapping

We used activity flow mapping – an empirical approach to model the propagation of distributed activity across brain regions in data – to evaluate cortical heterogeneity in distributed and localized processes. A core assumption of activity flow mapping is that the local task-evoked activity of a region is predicted by distributed neural processes (Cole et al., 2016). Here we tested whether there were differences in activity flow mapping predictions across cortical areas.

We estimated our resting-state FC weights using multiple linear regression, as was previously reported in (Cole et al., 2016). The benefit of using multiple linear regression to obtain the FC weights to a given target region y (relative to pairwise correlation) is that it partials out the time series of all other brain regions, while optimizing for prediction on y. Specifically, FC weights for a target region y was obtained by fitting the multiple linear regression model

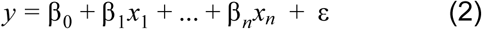

where the time series of all other regions {*x*_1_, …, *x*_*n*_} were used as regressors to predict region y, and coefficients {β_1_, …, β_*n*_ } corresponded to the FC weights used to predict region y’s task-evoked activation level. (Note that in the above equation n=359, since there are 360 parcels in the Glasser atlas.)

To obtain activity flow-mapped predictions for a target region for a given task, we mapped the task-evoked activations from all regions (excluding the target region) to the target region (Cole et al., 2016). Specifically, for a region y, the task-evoked activity prediction for a given task condition was defined as

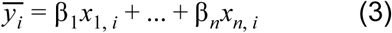

where 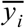 is the activity flow prediction for region y during task condition i, *x*_*n,i*_ is the true task-evoked activation for region *x* during task condition i, and β_*n*_ reflects the FC weight from region n to i obtained from equation 2.

Finally, to assess how well a region was predicted through the activity flow mapping procedure, we estimated the ‘activity flow mean absolute error’ (activity flow MAE) across all task conditions i=[1, …, 24]. Specifically, activity flow MAE for a region y was defined as

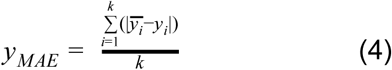

where *y*_*MAE*_ indicates the activity flow MAE for region y, *y*_*i*_ reflects the true task activation for region y during condition i, and k=24, which reflects the number of total task conditions (split among 7 tasks) in the HCP data set.

### Non-parametric statistical testing

All correlation-based statistical tests were performed using spatial autocorrelation-preserving permutation tests that generated random surrogate brain maps (Alexander-Bloch et al., 2018; Burt et al., 2018). We used the recently released BrainSMASH toolbox to generate 1000 surrogate brain maps for each variable of interest (e.g., task activation, task FC change, myelin, intrinsic timescale, and activity flow maps) (Burt et al., 2020). P-values were estimated from the null distribution of correlation values obtained by correlating each surrogate map with the variable of interest. We used Spearman’s rank correlation as our test statistic, though we obtained virtually identical results with Pearson’s correlation.

### Data and code availability

All code related to analyses in this study will be publicly released on GitHub. All data is publicly available through the Human Connectome Project (http://www.humanconnectomeproject.org) (Van Essen et al., 2013).

## Results

### Task-related activity and functional connectivity are differentiated across the cortical hierarchy

Task-related activity and FC are commonly used to characterize localized and distributed cognitive processes. Here we sought to evaluate whether regional differences in localized and distributed processes was related to macroscale cortical organization. Using task data collected from the Human Connectome Project (HCP) across 24 distinct task conditions, we directly compared how localized and distributed processes is differentiated across cortex by comparing the regional task activation strength and inter-region FC strength across multiple task conditions (Figure 1).

**Figure 1.**
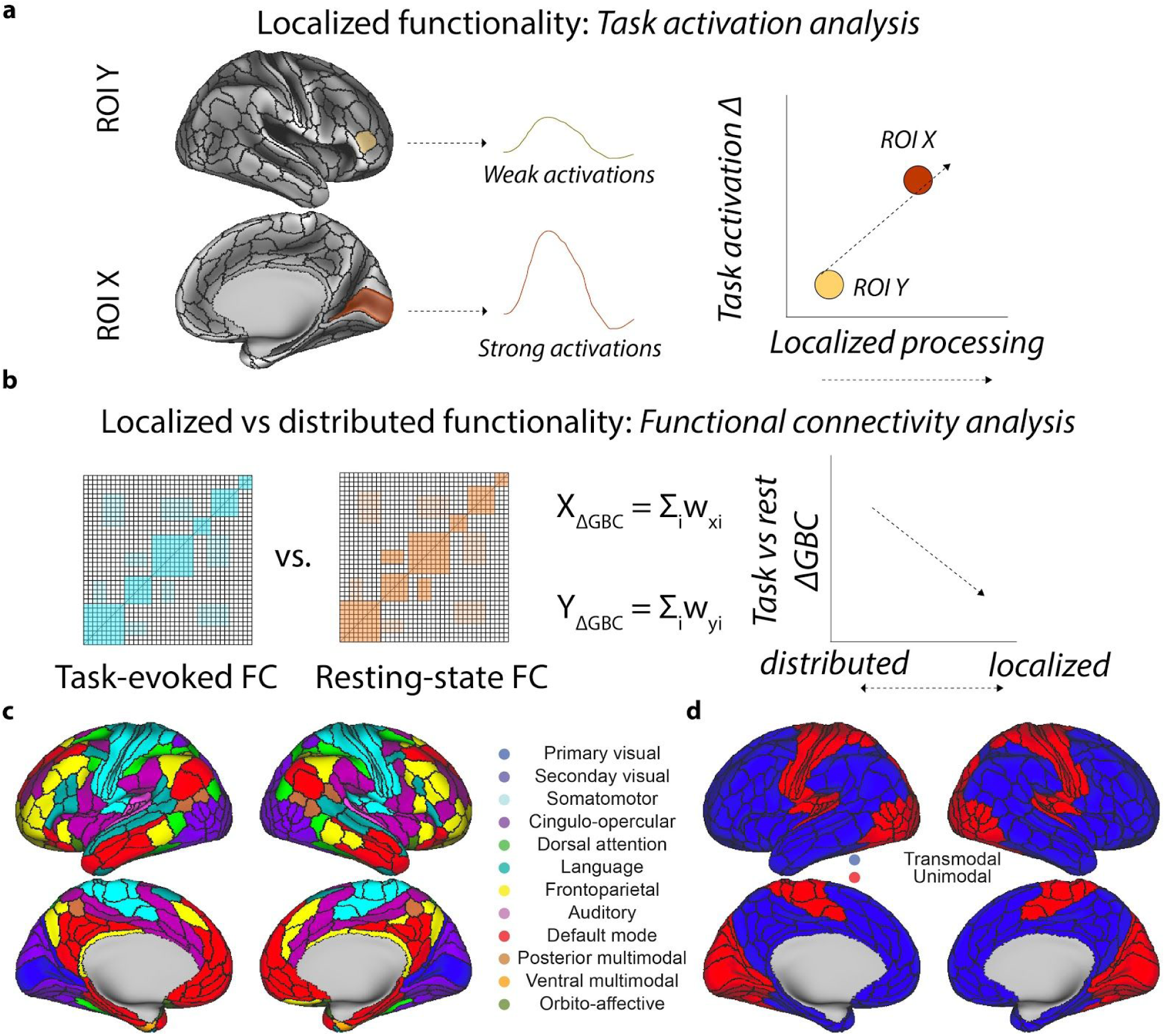
Data analysis schematic for assessing localized versus distributed processes during multiple task states. **a)** Characterizing localized functionality by estimating regional task activation changes. To identify the task activation change of each brain region, we estimated the task activation magnitude of each brain region across 24 task conditions. Localized processes were operationally defined as the magnitude of task activation change relative to baseline (see Methods). **b)** Characterizing a region’s distributed functionality by estimating the strength of its global FC strength relative to its resting-state FC. To identify the global FC change of a region, we compared the task-state global FC and compared it relative its resting-state global FC. Thus, a region’s reduced global FC during task states indicated that it was processing information in a more localized manner. **c,d)** To more simply compare localized and distributed processes across cortical areas, we mapped the previously described functional network assignments (Ji et al., 2019) into transmodal and unimodal networks. Unimodal networks included: primary and secondary visual networks, auditory network, and somatomotor network. Transmodal networks included all other networks.

To identify which regions were primarily involved in localized processes, we estimated the average magnitude (i.e., absolute value) of task-evoked activity across 24 task conditions for every parcel in the Glasser et al. 2016 atlas (Glasser et al., 2016). We found that unimodal regions had significantly higher task activation magnitudes as compared to transmodal regions, suggesting that unimodal areas respond more locally to tasks (Figure 2b,d; transmodal vs. unimodal, t(175) = −29.58, p<10e-69; replication set, t(175) = −27.07, p<10e-63). Congruent with increased local processes, we found that the FC of unimodal areas was also significantly reduced relative to transmodal areas during task states (Figure 2c,e; transmodal vs. unimodal, FC difference = 0.02, t(175) = 18.91, p<10e-43; replication set, FC difference = 0.03, t(175) = 20.63, p<10e-47).

**Figure 2.**
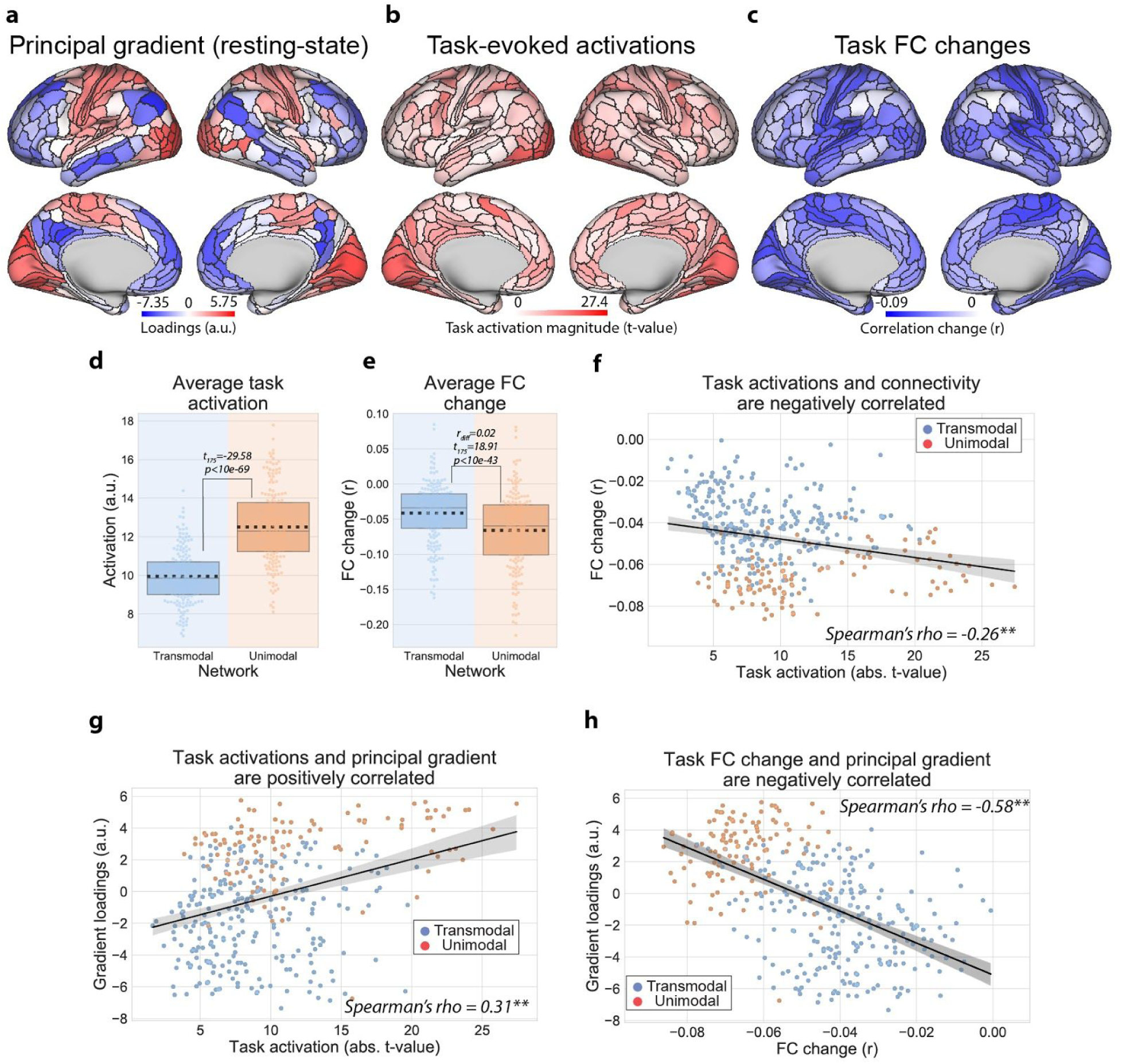
Dissociating localized versus distributed processes across the cortical hierarchy by estimating regional task activations and FC changes. **a)** The resting-state principal macroscale gradient (PG1) from Margulies and colleagues, which provides a spatial framework to characterize unimodal to transmodal activity (Margulies et al., 2016). **b)** Task activation magnitudes relative to baseline (absolute t-values), averaged across 24 task conditions. **c)** Averaged task FC changes for each region relative to resting-state FC. **d)** The task activation magnitudes averaged within transmodal and unimodal regions. Unimodal regions had significantly higher task activation magnitudes across multiple tasks relative to transmodal regions. Boxplots indicate the interquartile range of the distribution, dotted black line indicates the mean, grey line indicates the median, and the distribution is visualized using a swarm plot. **e)** Averaged task FC changes (relative to resting state) for transmodal and unimodal regions. In contrast to the task activation magnitude, unimodal regions significantly decreased their FC relative to transmodal regions. **f)** We also reproduced a result from our previous study (Ito et al., 2019a), demonstrating that regions with higher task-evoked activations decreased their FC more during task states. **g)** Task activation magnitudes were positively correlated with PG1. **h)** Task FC changes were negatively correlated with the PG1. All p-values (for correlation analyses) were estimated using a spatial autocorrelation-preserving permutation test to generate random surrogate brain maps (Burt et al., 2020). (*** = p<0.001, ** = p<0.01, * = p<0.05)

In a recent study, we found that the task-state FC of each region decreased (relative to resting state), and the magnitude of FC reduction was correlated with the magnitude of task activation increase (reproduced in Figure 2c). In a computational model, these FC reductions could be explained by the strengthening of a task-state attractor, which suppresses background spontaneous activity to help increase the fidelity of task signals (i.e., task-evoked activity) (Ito et al., 2019a). Importantly, these task-state correlation reductions are a phenomena that are commonly observed in electrophysiology data (Cohen and Kohn, 2011; Doiron et al., 2016), and have been proposed to reflect task-related information coding properties (Averbeck et al., 2006; Bartolo et al., 2020; Bejjanki et al., 2017; Zhang et al., 2019). Here we extended those results, and first reproduced the finding that regions that increased their task-evoked activations tended to decrease their task-state FC (Figure 2f) (Ito et al., 2019a). This illustrated that localized (local task activations) and distributed processes (region’s average FC) were negatively correlated across regions (Spearman’s rank correlation (r_s_) = −0.26, bootstrapped 95% confidence intervals (CI_95_) = (−0.36, −0.16), p=0.004; replication set, r_s_ = −0.27, CI_95_ = (−0.36, −0.17), p=0.002). We note that this negative association also held when thresholding the FC matrices to include only positive FC values (Supplementary Figure 4). (All p-values for correlation-based statistics for brain maps were obtained by using a spatial autocorrelation-preserving permutation test that generates random surrogate brain maps (see Methods) (Burt et al., 2020).)

Importantly, the topographic changes in task activations and task-state FC were associated with the recently-published resting-state macroscale principal gradient (PG1), which provides a spatial framework for characterizing unimodal to transmodal activity (Margulies et al., 2016). Specifically, topographic changes in task activation magnitudes were positively correlated with the PG1 map (Figure 2g; r_s_ = 0.31, CI_95_ = (0.22, 0.40), p = 0.004; replication set, r_s_ = 0.28, CI_95_ = (0.17, 0.38), p = 0.006), while task-state FC changes were negatively correlated with the PG1 map (Figure 2h; r_s_ = −0.58, CI_95_ = (−0.63, −0.51), p<0.001; replication set, r_s_ = −0.57, CI_95_ = (−0.63, −0.51), p<0.001). These findings illustrated a dissociation of localized and distributed processes across transmodal and unimodal networks, indicating unimodal networks respond to tasks more locally, while transmodal networks respond to tasks more distributedly. Moreover, these changes in task-related activity and FC were associated with the principal macroscale gradient previously observed from resting-state fMRI (Margulies et al., 2016), which suggests a potential organizing principle underlying spontaneous and task-evoked states.

### Task-state functional cortical heterogeneity is related to intrinsic timescales during resting state

The negative association between task-related activations and FC changes illustrates the relationship between distributed and localized processes within the cortex. However, it is unclear why such a functional dichotomy exists. A previous study showed that the intrinsic timescales across cortical areas follow anatomical connectivity maps in non-human primates, suggesting that anatomical wiring organization and functional timescale hierarchies are closely related (Murray et al., 2014). Moreover, that same study indicated lower-order cortical areas tend to operate at fast timescales, potentially supporting stimulus-/task-locked activity. However, conclusions from Murray et al. (2014) were based on non-human primate electrophysiology data. Thus, we first sought to identify an intrinsic timescale hierarchy in human fMRI data, and additionally hypothesized that the dissociation of regional task activations and FC changes were associated with differences in a brain region’s intrinsic timescale.

We measured the intrinsic timescale of brain regions using resting-state fMRI. We fitted a nonlinear exponential decay function to the empirically calculated autocorrelation function (100 timepoint lag), and used the decay rate (τ) as the intrinsic timescale (Murray et al., 2014). This intrinsic timescale represents the rate of decay (of the autocorrelation function) to 0. (Results were consistent when fitting to shorter lags, including 40 and 50 time points.) This meant that regions with a larger decay parameter (τ) had a slower decay rate. (An alternative interpretation is that regions with a longer timescale (larger τ) have more autocorrelation in the time series and slower temporal fluctuations.) We found that transmodal regions had significantly slower intrinsic timescales than unimodal regions (Figure 3c; τ difference = 0.99, t_175_=19.33, p<10e-44; replication set τ difference = 0.96, t_175_=22.12, p<10e-51). These findings corroborate a previous study in non-human primate electrophysiology reporting that lower-order cortical areas tend to operate at fast timescales, while higher-order areas tend to operate at slow timescales (Murray et al., 2014).

**Figure 3.**
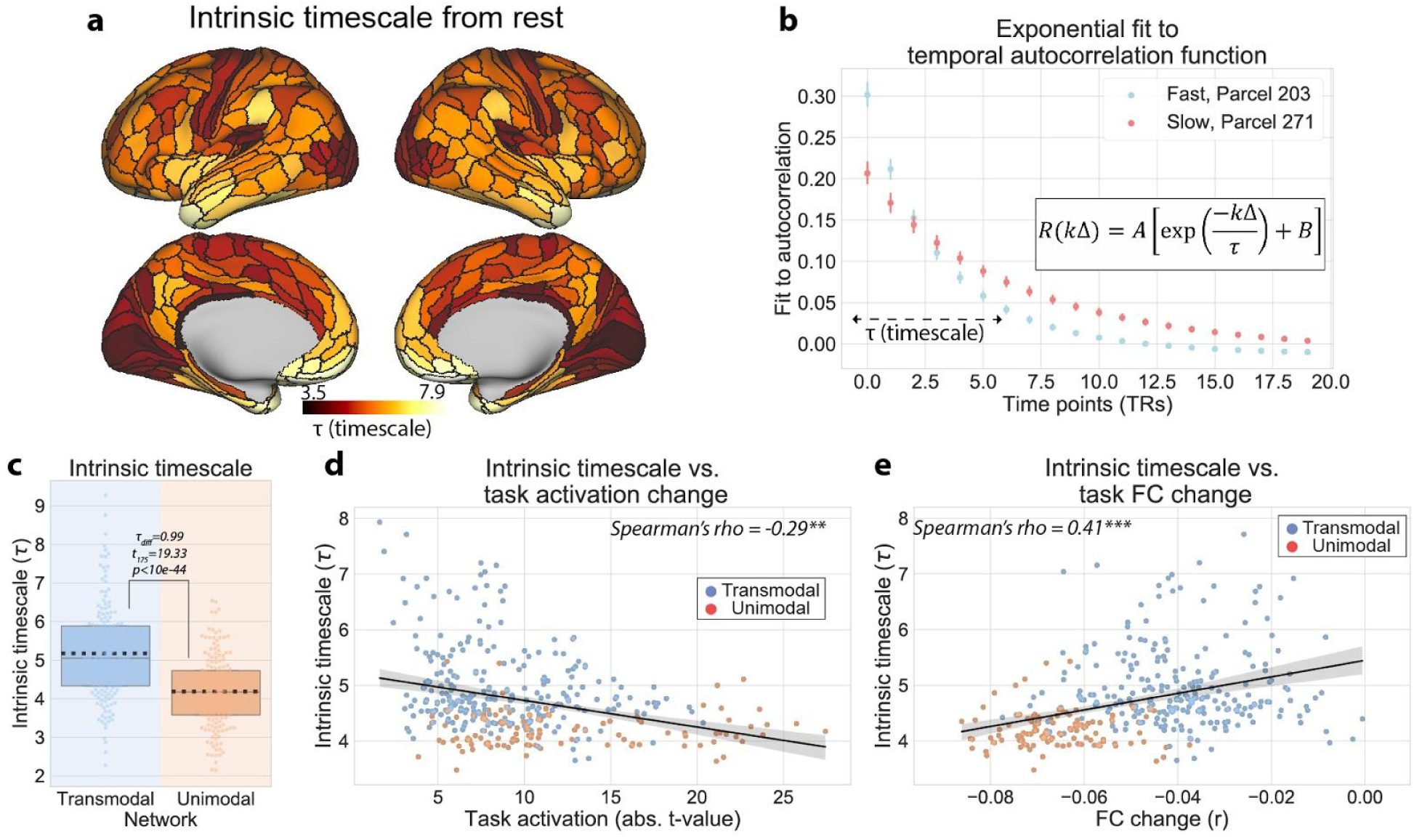
Hierarchy of intrinsic timescales estimated during resting-state fMRI explains regional differences in task activations and FC. **a)** The intrinsic timescale for each cortical region. We estimated the intrinsic timescale of each region by fitting a 3-parameter exponential decay function to the autocorrelation function obtained during resting-state fMRI (Murray et al., 2014). **b)** The estimated exponential decay functions for two example regions with fast (blue) and slow (red) timescales. Fits were estimated for each subject separately. Error bars denote the 95% confidence interval (across subjects). **c)** The intrinsic timescale (i.e., the rate of decay) was significantly greater for transmodal regions relative to unimodal regions. Boxplots indicate the interquartile range of the distribution, dotted black line indicates the mean, grey line indicates the median, and the distribution is visualized using a swarm plot. **d)** Across the cortical hierarchy, the intrinsic timescale was negatively correlated with task activation magnitude across multiple tasks, consistent with the notion that regions with fast timescales respond in a stimulus/task-locked manner. **e)** In contrast, the intrinsic timescale was positively correlated with the task-state FC change, consistent with the hypothesis that regions with slow timescales have a larger temporal receptive field and can integrate information from lower-order cortical areas. All p-values (for correlation analyses) were estimated using a spatial autocorrelation-preserving permutation test to generate random surrogate brain maps. (*** = p<0.001, ** = p<0.01, * = p<0.05)

Other studies demonstrated that higher-order cortical regions integrate information at slower timescales relative to lower-order regions during naturalistic/continuous stimuli (Baldassano et al., 2017; Hasson et al., 2008; Honey et al., 2012). It is thought that regions operating at slower intrinsic timescales are more likely to integrate information from other regions, similar to the feedforward and compressive temporal summation principles observed in visual cortex (Cocchi et al., 2016; Zhou et al., 2017). In contrast, regions operating at fast timescales should respond in a more stimulus-/task-locked manner. Thus, we hypothesized that regions with faster intrinsic timescales are more likely to have higher task activation magnitudes (given their likelihood of having more stimulus-/task-locked neural responses), while simultaneously reducing their FC strength due to a lesser ability to temporally integrate information from other brain regions. Indeed, we found that across the cortical hierarchy, regions with faster timescales (i.e., smaller τ values) had larger task activation magnitudes (Figure 3d; r_s_ = −0.29, CI_95_ = (−0.38, −0.19), p=0.002; replication set, r_s_ = −0.29, CI_95_ = (−0.39, −0.19), p<0.001). In addition, we found that regions with faster timescales had also reduced their task-state FC more (Figure 3e; r_s_ = 0.41, CI_95_ = (0.32, 0.49), p<0.001; replication set, r_s_ = 0.43, CI_95_ = (0.34, 0.52), p<0.001).

### Task-state functional cortical heterogeneity is related to local myelin density

The above results show that hierarchical differences of intrinsic timescales estimated during resting state correspond to differences in task-related activation and FC changes. However, previous reports also indicate that such hierarchical cortical heterogeneity may be due to structural, genetic, and synaptic differences (Burt et al., 2018; Demirtaş et al., 2019; Huntenburg et al., 2018; Vázquez-Rodríguez et al., 2019; Wang, 2020). Thus, we sought to demonstrate that the hierarchy of intrinsic timescales in human fMRI is related to changes in structural differences (i.e., myelination content), while extending these associations to incorporate the cortical heterogeneity in localized and distributed processing.

Theoretical work has demonstrated that strong local coupling of excitatory-inhibitory (E-I) connectivity generate fast neural dynamics (Hennequin et al., 2018, 2017; Ito et al., 2019a; Lombardi et al., 2017; Wang et al., 2019). This has been corroborated in recent empirical studies, where cortical heterogeneity of local E-I coupling (which was highly similar to regional myelin content) produced simulated neural dynamics that closely matched large-scale human fMRI data (Demirtaş et al., 2019). Thus, we hypothesized that cortical heterogeneity in myelin content (Figure 4a) would be related to both the hierarchy of intrinsic timescales and differences in localized and distributed processes.

**Figure 4.**
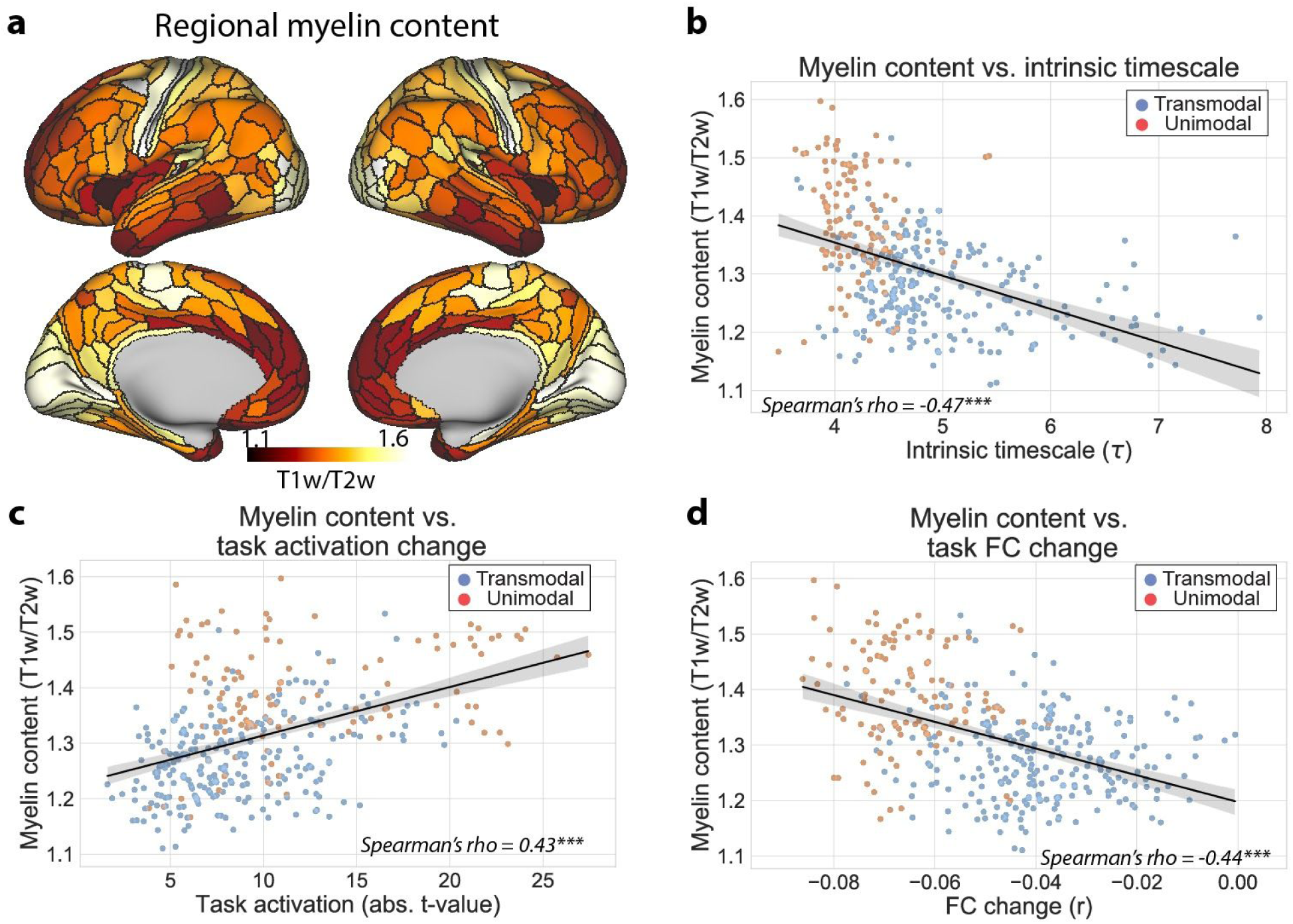
Intrinsic and task-state differences in hierarchical cortical organization are associated to local myelin density. **a)** Cortical myelin content within each parcel estimated from a T1w/T2w contrast map (Burt et al., 2018). **b)** Across cortical regions, myelin content and the intrinsic timescale are negatively related, suggesting that lower-order brain regions operate at faster intrinsic timescales. **c)** Across cortical regions, myelin content is positively correlated with the magnitude of task-evoked activations, suggesting that lower-order brain regions tend to have higher task-evoked activations (consistent with stimulus-locked activity). **d)** Across cortical regions, myelin content is correlated with task-state FC decreases, suggesting that higher-order brain regions change their task FC strength less (consistent with information integration with other brain regions). All p-values were estimated using a spatial autocorrelation-preserving permutation test to generate random surrogate brain maps. (*** = p<0.001, ** = p<0.01, * = p<0.05)

We found a negative association between regional myelin content and the intrinsic timescale across cortical regions (Figure 4b; r_s_ = −0.47, CI_95_ = (−0.55, −0.37), p<0.001, replication set, r_s_ = −0.49, CI_95_ = (−0.57, −0.40), p<0.001). This suggests that regions with higher myelin content (i.e., local connectivity), such as unimodal areas, operate at faster intrinsic timescales. In addition, we found that regional myelin content was positively correlated with the magnitude of task activations (Figure 4c; r_s_ = 0.43, CI_95_ = (0.34, 0.51), p<0.001; replication set, r_s_ = 0.41, CI_95_ = (0.32, 0.50), p<0.001), while negatively correlated with the reduction of a region’s average task FC (Figure 4d; r_s_ = −0.44, CI_95_ = (−0.51, −0.35), p<0.001; replication set, r_s_ = −0.46, CI_95_ = (−0.54, −0.37), p<10e-19). Together, these results suggest that unimodal regions have more local coupling (i.e., myelin content), driving faster intrinsic timescales during rest and task-locked neural responses during tasks. In contrast, our results suggest that transmodal regions have less local E-I coupling, facilitating slower intrinsic timescales (i.e., wider temporal receptive field) and weaker task-state FC changes.

### Dissociation of activity and functional connectivity across cortex is not dependent on task-based regression

The previous results used a standard task general linear model (GLM) to estimate task activations while using the residual time series after removing the main effect of task for FC analyses (Cole et al., 2019). However, it is possible that the estimated activity from a standard GLM coefficient may not appropriately capture activity associated with the task (e.g., due to linear approximation of a task GLM). This is because standard task GLM coefficients are extracted from a stationary task block design convolved with a canonical hemodynamic response function (Friston et al., 1994). Similarly, the FC estimates we used capture the timepoint-to-timepoint variance that is left over after task regression. However, other approaches have illustrated that FC can also be obtained without using the residual time series by estimating the trial-to-trial (or block-to-block) variance of task-evoked activation levels (Ito et al., 2019a; Rissman et al., 2004). Thus, we sought to demonstrate that the negative relationship between mean task-related activity and FC observed above was not dependent on any task GLM (or FIR) modeling.

Using preprocessed fMRI time series (without applying any task GLM or FIR model), we obtained the peak activation value within each task block relative to baseline for each block across the 7 HCP tasks (see Methods). These values represented the peak activation value for each region within each task block, independent of whether the activity was sustained across the entire block. To obtain the task activation magnitude for a cortical region, we averaged the activation peaks across all blocks for every task condition relative to baseline. To obtain the task-state FC change for a pair of brain regions, we computed the correlation of peak activations across all blocks (rather than time points) between pairs of brain regions and evaluated the FC change relative to resting-state (see Methods). The estimated task FC is conceptually similar to the beta series FC approach (Rissman et al., 2004), which captures event-to-event variance rather than timepoint-to-timepoint variance. (However, rather than using the estimated beta coefficient from a task GLM, we use the peak BOLD value at each event.)

We found that all associations (both positive and negative) between activity, FC change, intrinsic timescales, and myelin content were replicated using the peak activation approach (Figure 5b). Moreover, the associations observed with the peak activation approach were higher than the standard task GLM approach to measure task activations and FC change (Figure 5a). These results demonstrate that the dissociation of task activations and task FC is independent of task regression methods. Specifically, task FC can be dissociated from task activations at both timepoint-to-timepoint and block-to-block temporal scales.

**Figure 5.**
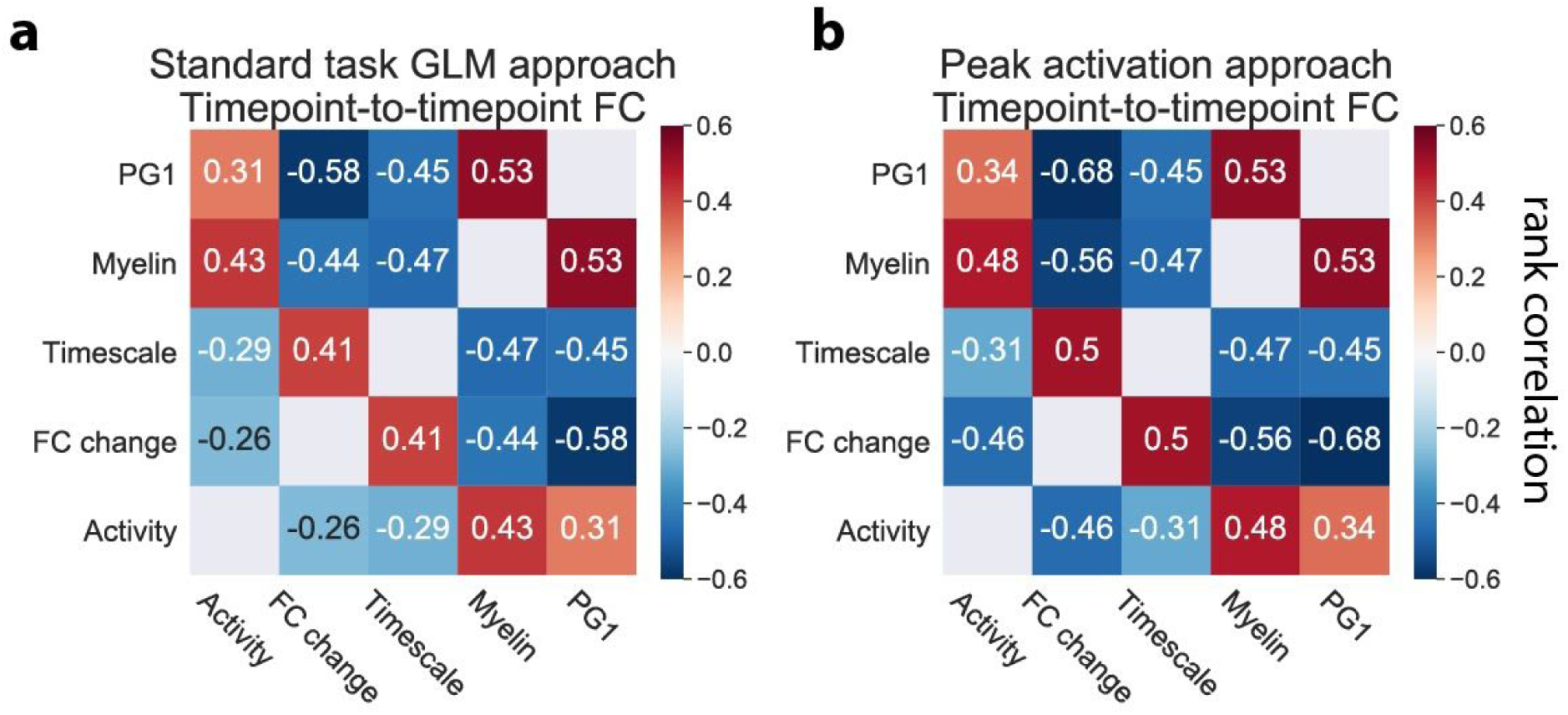
Summary of positive and negative associations between the resting-state principal gradient, task activations, task FC change, intrinsic timescales, and myelin content using the standard task GLM and the peak (block) activation approaches. (Figures for the replication cohort are in Supplementary Figure 1.) **a)** The standard task GLM approach for the exploratory cohort. We use standard task GLM modeling to estimate activation coefficients for each brain region, and FIR task modeling to remove the mean task-evoked response prior to computing task FC. Note that all measures reported in this study were strongly associated with the principal resting-state gradient (PG1), which was hypothesized to reflect hierarchical organization in the brain (Margulies et al., 2016). **b)** The peak activation approach for the exploratory cohort. We estimate the peak activation magnitude at each block (across all blocks) without task regression. Task activations are estimated by averaging peak magnitudes across all blocks for each brain region. Task FC estimates are obtained by correlating block-to-block variance (using peak magnitudes) between all pairs of brain regions. Positive and negative association strengths are typically stronger using the peak activation approach. All correlations were found to be statistically significant using an FDR-corrected p-value of p<0.01. All p-values were estimated using a spatial autocorrelation-preserving permutation test to generate random surrogate brain maps.

### Improved activity flow mapping predictions of transmodal areas due to distributed processes

Previous work from our group has demonstrated that the task-evoked activity of distributed brain regions can be predicted by modeling ‘activity flow’ processes over functional weights estimated from resting-state fMRI (Cole et al., 2016; Ito et al., 2019b, 2017). These activity flow processes are modeled by predicting a brain region’s task-evoked activation level by summing the task-evoked activations of all other brain regions weighted by their resting-state FC weights to the predicted region (Figure 6a; see Methods). Specifically, we empirically estimate FC weights (from resting-state data) then simulate the propagation of activity of time-resolved task-evoked activations over those FC weights to predict activity in each brain region. The simulation of ‘activity flow’ is equivalent to simulating artificial neural network computations (formally called the propagation rule) (Rumelhart et al., 1986). Instead of using correlation to estimate FC weights, we employ multiple linear regression, since it conditions on all other brain regions when estimating the weights between any two brain regions, reducing causal confounds (Cole et al., 2016).

**Figure 6.**
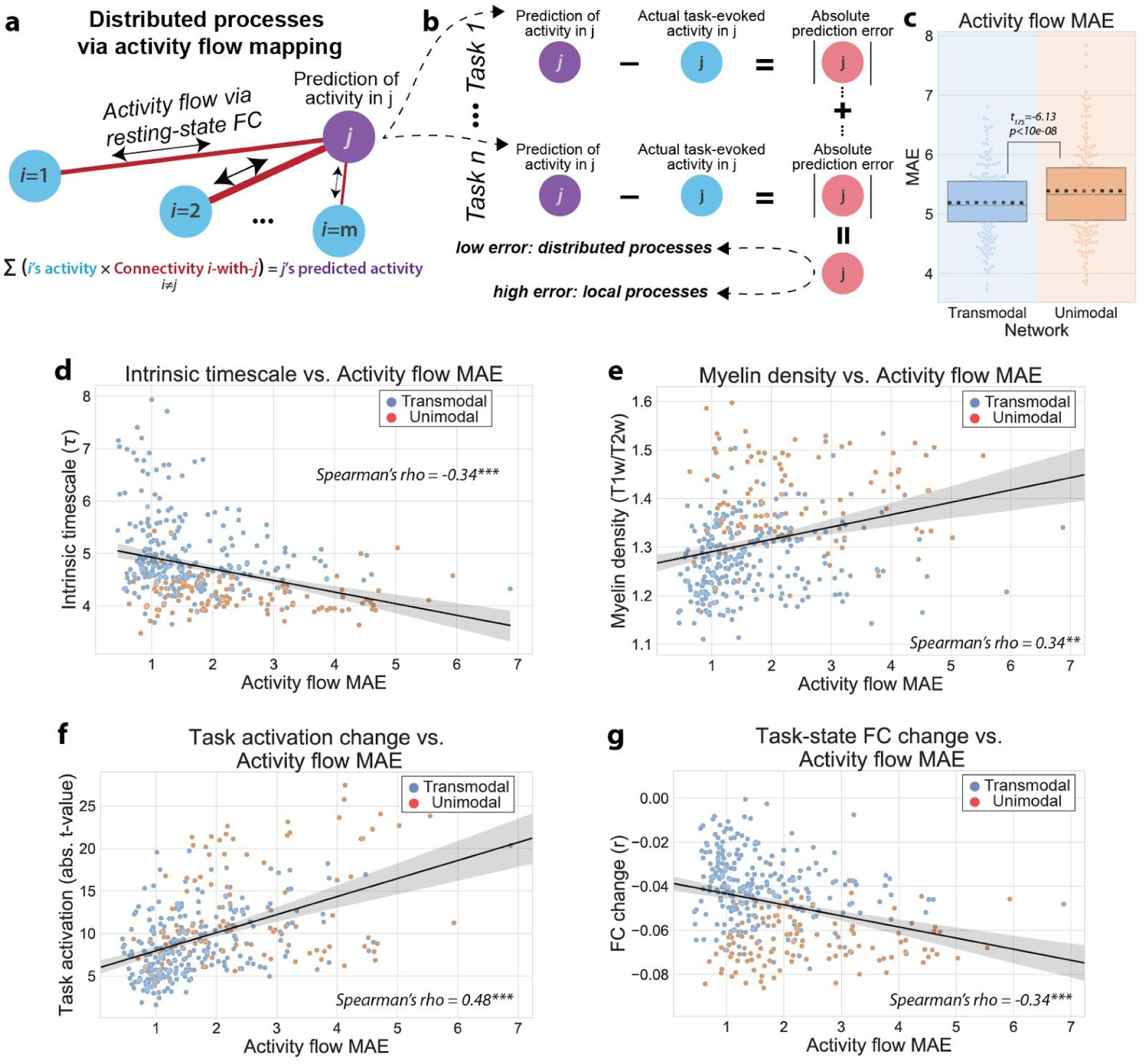
Better prediction of task-evoked activations for transmodal regions than unimodal regions via activity flow mapping. **a)** The activity flow mapping algorithm, which was originally derived from connectionist principles (Cole et al., 2016; Ito et al., 2019b). Briefly, the task-evoked activation of a brain region j can be predicted by summing the task-evoked activations of all other brain regions weighted by their FC weights with region j. A core assumption of this algorithm is that the task-evoked activity of region j is generated from a distributed process, rather than from a local (or internal) process. **b)** To evaluate whether some brain regions are better predicted via activity flow mapping, we can characterize the mean absolute error of the activity flow predictions (i.e., ‘activity flow MAE’) across task conditions. This evaluates the mean absolute error of the activity flow mapping algorithm for every brain region. **c)** We find that transmodal regions have significantly lower activity flow MAE relative to unimodal regions. Boxplots indicate the interquartile range of the distribution, dotted black line indicates the mean, grey line indicates the median, and the distribution is visualized using a swarm plot. **d)** We found a negative association between the intrinsic timescale of regions and activity flow MAE (i.e., slower intrinsic timescales have better activity flow predictions, consistent with the view that a wider temporal receptive field facilitates better information integration from different regions). **e)** We found a positive association between the myelin content of regions and activity flow MAE. This is consistent with the notion that lower-order cortical regions process information more locally. **f,g)** Across cortical regions, activity flow MAE was positively/negatively associated with task activation magnitude/FC change. (*** = p<0.0001, ** = p<0.01, * = p<0.05)

However, a core assumption of activity flow mapping is that the activity of a brain region is the result of distributed processes, since its activity can be predicted from the activity of other brain regions. However, the present results suggest that there is hierarchical heterogeneity in localized and distributed processes, as evidenced by differences in regional task activations and FC changes (Figure 2). Thus, consistent with the dichotomy of localized and distributed functional processes, we hypothesized that regions more involved in distributed processes (e.g., transmodal regions) would be better predicted by activity flow mapping relative to regions with more localized processes (e.g., unimodal regions). This implies that the localized processes in unimodal regions are not as well explained by distributed activations.

We applied the activity flow algorithm to predict the task-evoked activation of every brain for every task condition. To assess how well a target brain region could be predicted by activity flow from other distributed brain regions, we computed the absolute value of the error between the predicted and actual task-evoked activity for each task condition (activity flow mean absolute error (MAE)). We then averaged across all task conditions (i.e., the mean absolute error of activity flow predictions), providing an estimate of the activity flow MAE for each brain region (Figure 6b). Thus, activity flow MAE was a measure of how well a target region’s activity could be predicted as a function of distributed processes. (Lower activity flow MAE corresponded to better prediction.) We found that the average activity flow MAE was significantly lower in transmodal regions relative to unimodal regions, suggesting that transmodal regions were better predicted by activity flow mapping (Figure 6c; t(175)=-6.13, p<10e-08; replication set, t(175)=-3.27, p=0.001). We then correlated task-evoked activity flow MAE with two of the previous task-free estimates that describe hierarchical cortical organization: the intrinsic timescale and myelin content. Indeed, we found that the intrinsic timescale was negatively correlated with the amount of activity flow MAE across cortical areas (Figure 6d; r_s_ = −0.34, CI_95_ = (−0.43, −0.24), p<0.001; replication set, r_s_ = −0.38, CI_95_ = (−0.47, −0.28), p<0.001), suggesting that it was harder to predict the task activations of regions with faster operating timescales. Similarly, we found that myelin content was negatively correlated with activity flow MAE (Figure 6e; r_s_ = 0.34, CI_95_ = (0.25, 0.43), p=0.001; replication set, r_s_ = 0.31, CI_95_ = (0.21, 0.42), p=0.001), indicating that regions with more local coupling (higher myelination) are harder to predict via distributed FC weights. We also found that activity flow MAE in each region was positively correlated with its true task activation magnitude (Figure 6f; r_s_ = 0.48, CI_95_ = (0.39, 0.56), p<0.001; replication set, r_s_ = 0.42, CI_95_ = (0.32, 0.51), p<0.001), while simultaneously being negatively correlated with the task-state FC change in each region (Figure 6g; r_s_ = −0.34, CI_95_ = (−0.43, −0.24), p<0.001; replication set, r_s_ = −0.32, CI_95_ = (−0.41, −0.22), p<0.001). Together, these results are congruent with the overall hypothesis that unimodal regions reflect more local processes, while transmodal regions reflect more distributed processes.

## Discussion

Our results provide evidence for a cortical hierarchy of localized and distributed processes revealed by differences in task activations, task FC changes, intrinsic timescales, and myelin content. We found that across multiple tasks, regions with high-levels of stimulus-/task-locked activity tended to reduce their global FC, suggesting they processed information more locally. Specifically, unimodal regions tended to activate and reduce their FC during task states relative to transmodal regions, consistent with the notion that unimodal regions respond more locally, while transmodal regions respond distributedly (Cole et al., 2013; Huntenburg et al., 2018). Moreover, these differences were linked to several intrinsic (task-free) properties of macroscale cortical organization: hierarchical timescale organization and cortical myelination. Since fMRI measures the blood-oxygen-level-dependent (BOLD) signal, which is indirectly related to metabolic and neural activity (Logothetis et al., 2001; Ma et al., 2016), our findings suggest that cortical heterogeneity of intrinsic properties may drive differences in localized and distributed neural and/or metabolic processes.

We found a hierarchy of intrinsic timescales from unimodal to transmodal regions operating at fast to slow timescales, consistent with previous reports (Murray et al., 2014). Moreover, we found that the hierarchy of timescales was negatively correlated with local processes (task activation magnitudes) and positively correlated with distributed processes (task FC change). This indicated that regions with faster intrinsic timescales (primarily unimodal regions) activated in a stimulus-/task-locked manner and decreased their FC more during task.

To assess which regions processed information more distributedly, we used our previously-developed activity flow mapping approach, which assumes local neural activity can be predicted from the propagation of activity flow from distributed brain areas (Cole et al., 2016). We found that transmodal regions yielded better activity flow predictions across multiple tasks, indicating that the activity of transmodal regions result from distributed processing. Importantly, the accuracy of task-evoked activity predictions was correlated with the hierarchy of the intrinsic timescale organization, suggesting that regions with slower timescales were better predicted by distributed activity flow processes. These findings are consistent with the notion that regions with slower timescales have a wider temporal receptive field to integrate information from other brain regions (Baldassano et al., 2017; Cocchi et al., 2016; Hasson et al., 2008; Honey et al., 2012).

We found that regional myelin content was correlated with the intrinsic timescale (negatively), task activation magnitudes (positively), and task FC changes (negatively). This is consistent with previous studies suggesting an anatomical basis for hierarchical cortical functionality in models predicting resting-state functional network organization (Burt et al., 2018; Demirtaş et al., 2019; Wang et al., 2019) and studies reporting gradients of structure-function connectivity tethering (Baum et al., 2020; Vázquez-Rodríguez et al., 2019). Our results extend those previous findings to suggest an anatomical basis for the hierarchy of timescale organization as well as differences in local and distributed processing during task states. Specifically, regions with higher white matter content tended to operate at faster timescales and process information more locally.

Regional myelin content has recently been linked to anatomical hierarchies and cortical gradients of gene expression, indicating a potential link between microscale anatomy and macroscale functional organization (Burt et al., 2018). In recent work, regional myelin content was correlated with the degree of local (regional) E-I coupling parameters in a whole-brain biophysical network model that was parameterized to reproduce resting-state dynamics (Demirtaş et al., 2019). Thus, myelin content appears to reflect the degree of local E-I coupling (Wang, 2020). Independently, other theoretical work showed that changes to local E-I coupling directly influences the timescale and frequency characteristics of neural activity, with faster timescales associated with stronger inhibitory feedback (i.e., increased E-I coupling) (Lombardi et al., 2017). This finding provided evidence for a hypothesis that links myelin content and intrinsic timescales: Regional differences in myelin content reflect differences in E-I coupling, which in turn may govern intrinsic timescale properties. Our results provide some evidence for this hypothesis, finding that higher myelin content was associated with faster timescales. However, our findings were only associational, and as recently discussed, future work will need to directly link regional myelin content, E-I coupling, and intrinsic timescale behavior (Wang, 2020).

A recent study from our group illustrated that task-related activity (task signal) and background spontaneous activity (neural noise) can be effectively dissociated when removing the main effect of task from task time series (Ito et al., 2019a). The separation of task ‘signal’ from neural ‘noise’ enhances the interpretation of ongoing cognitive processes associated during tasks, and has been the standard approach in electrophysiology for decades (referred to as signal and noise correlations, respectively) (Aertsen et al., 1989; Cohen and Kohn, 2011). (It should be noted that the term noise correlations does not imply a lack of neural information in a time series, but rather the isolation of information in relation to the task.) Under this interpretation, our results suggest that regions that tend to activate during task states suppress background neural noise. The suppression of background correlated noise supports increased fidelity of the task signal (i.e., mean task activity), consistent with previous reports in the electrophysiology literature (Baria et al., 2017; Churchland et al., 2011, 2012; Cohen and Kohn, 2011).

There is a concern that separating the main task effect from the time series may artificially induce a negative association of task activity and task FC. However, removing the main effect of task would not artificially reduce the correlation of distributed spontaneous activity because signal and noise are statistically orthogonal sources of a data distribution (MacKay, 2003). Importantly, failure to remove the task signal prior to estimating task FC would conflate signal and noise correlations, weakening the strength of possible inferences (Cole et al., 2019; Reid et al., 2019). Yet independent of these concerns, we demonstrated that task activity and task FC can be dissociated without using a task GLM or FIR model to separate the main effect of task from the underlying time series. This approach isolates block-to-block/event-to-event variance by estimating the peak BOLD activation within each task block. The mean task activation magnitude is then captured by the average across blocks, while the task FC is captured by correlating the peaks between brain regions (across blocks). (This latter approach is similar to computing task FC using a beta series correlation (Rissman et al., 2004).)

In conclusion, we provide evidence that cortical differences in task-related activity and FC dynamics are differentially related to the hierarchy of intrinsic cortical organization. Overall, we found that unimodal regions process information more locally, while transmodal regions process information more distributedly. We note that the negative relationship between local and distributed processes did not have to be true; it could have been the case that regions with high amounts of local activity actually increased their distributed interactions during tasks, suggesting the existence of hub nodes that respond both locally and distributedly. But in contrast to this alternative hypothesis, our findings are consistent with the notion that transmodal regions integrate information from lower-order regions due to a wider temporal receptive field, while unimodal regions respond to tasks in a stimulus-/task-locked manner. Finally, these differences in localized and distributed were related to differences in the anatomical hierarchy as measured by cortical myelin content. We expect these findings to spur additional investigations into characterizing hierarchical cortical function during resting and task states.

## Acknowledgements

We thank John D. Murray for providing access to the group averaged T1w/T2w (myelin) map. The authors acknowledge support by the US National Institutes of Health under awards R01 AG055556 and R01 MH109520. The data were provided by the Human Connectome Project, WU-Minn Consortium (Principal Investigators: David Van Essen and Kamil Ugurbil; 1U54MH091657) funded by the 16 NIH Institutes and Centers that support the NIH Blueprint for Neuroscience Research; and by the McDonnell Center for Systems Neuroscience at Washington University. The content is solely the responsibility of the authors and does not necessarily represent the official views of any of the funding agencies.

## Supplementary Figures

**Supplementary Figure 1.**
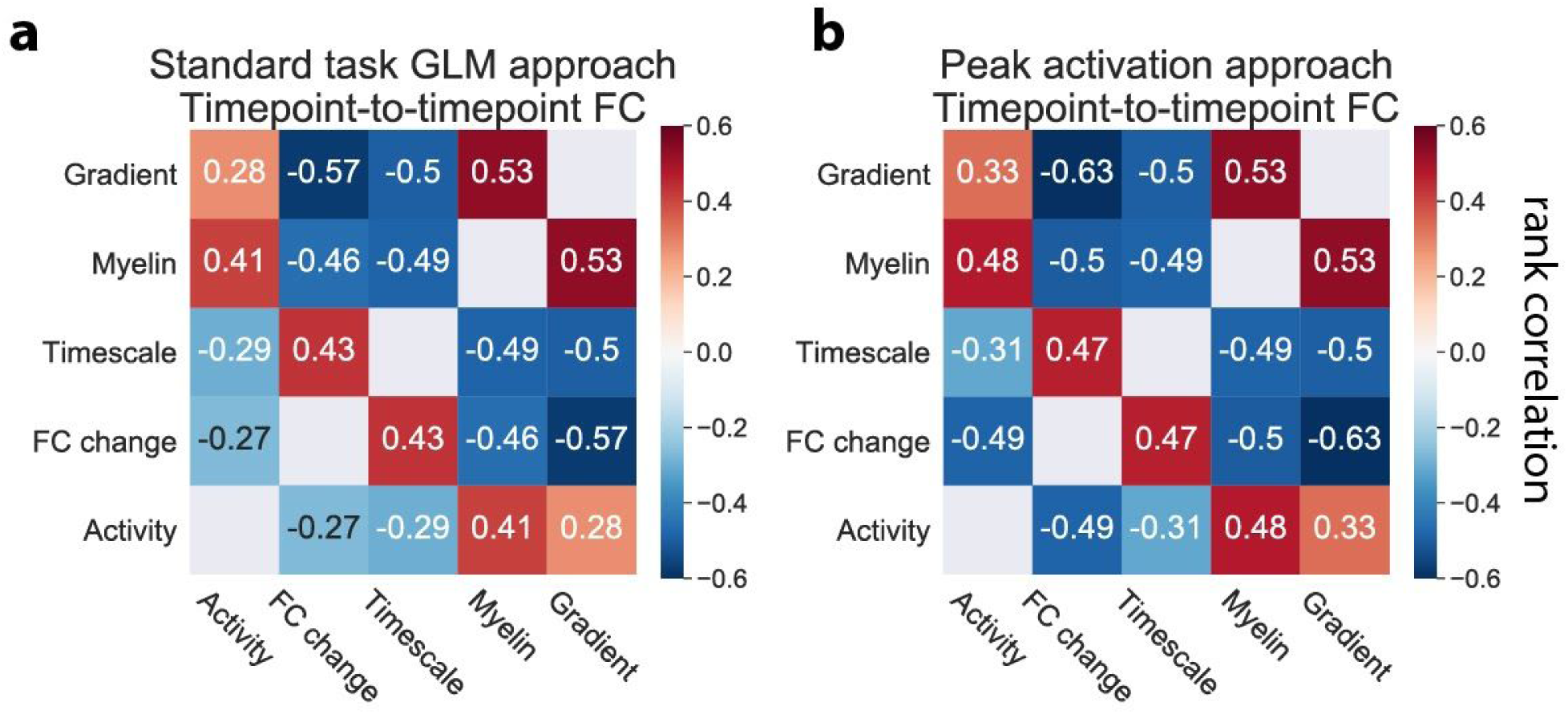
Summary of positive and negative associations (on the replication cohort) between the resting-state principal gradient, task activations, task FC change, intrinsic timescales, and myelin content using the standard task GLM and the peak (block) activation approaches. **a)** The standard task GLM approach for the replication cohort. We use standard task GLM modeling to estimate activation coefficients for each brain region, and FIR task modeling to remove the mean task-evoked response prior to computing task FC. **b)** The peak activation approach for replication. We estimate the peak activation magnitude at each block (across all blocks) without task regression. Task activations are estimated by averaging peak magnitudes across all blocks for each brain region. Task FC estimates are obtained by correlating block-to-block variance (using peak magnitudes) between all pairs of brain regions. Positive and negative association strengths are typically stronger using the peak activation approach. All correlations were found to be statistically significant using an FDR-corrected p-value of p<0.01. All p-values were estimated using a spatial autocorrelation-preserving permutation test to generate random surrogate brain maps.

**Supplementary Figure 2.**
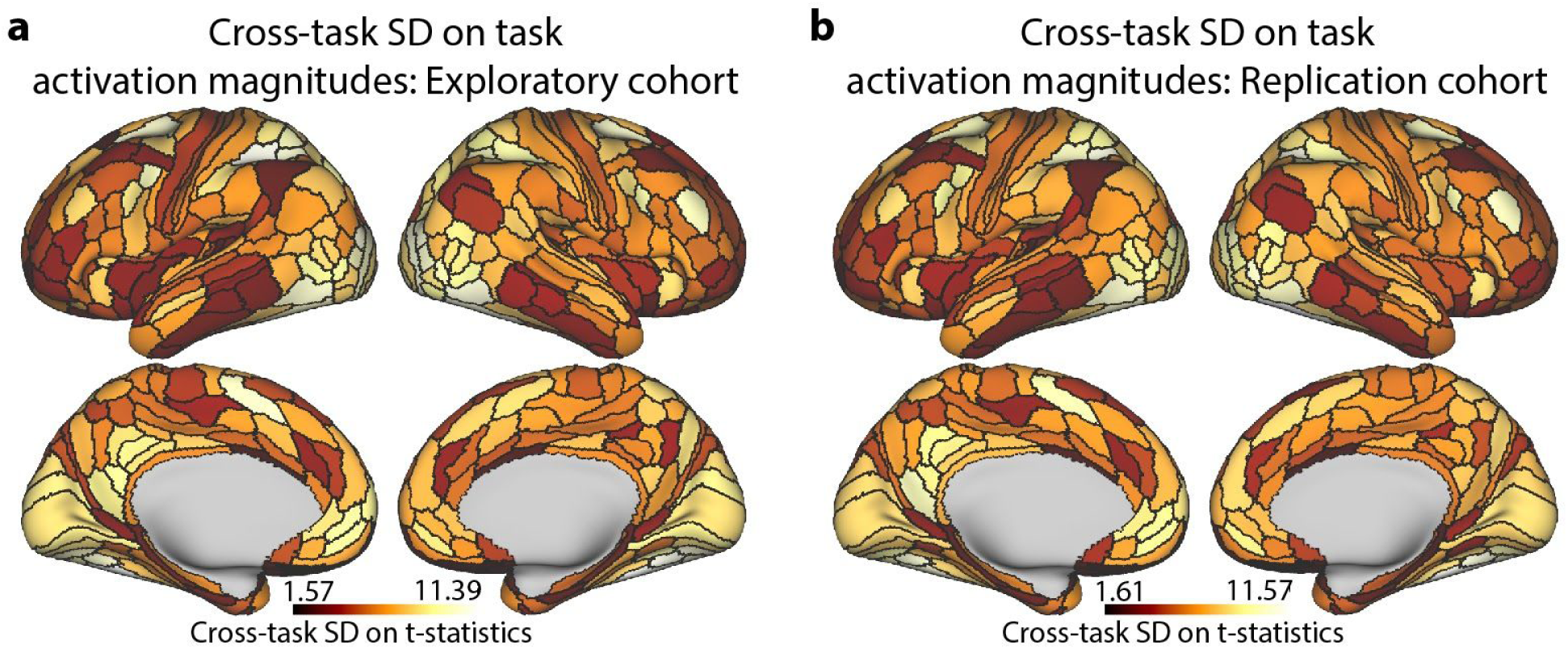
The standard deviation (across tasks) of task activation magnitudes at each parcel. **a)** Cross-task SD (on 24 task conditions) on the task activation magnitudes (absolute value of t-statistic) for the exploratory cohort. **b)** Same as **a**, but for the replication cohort.

**Supplementary Figure 3.**
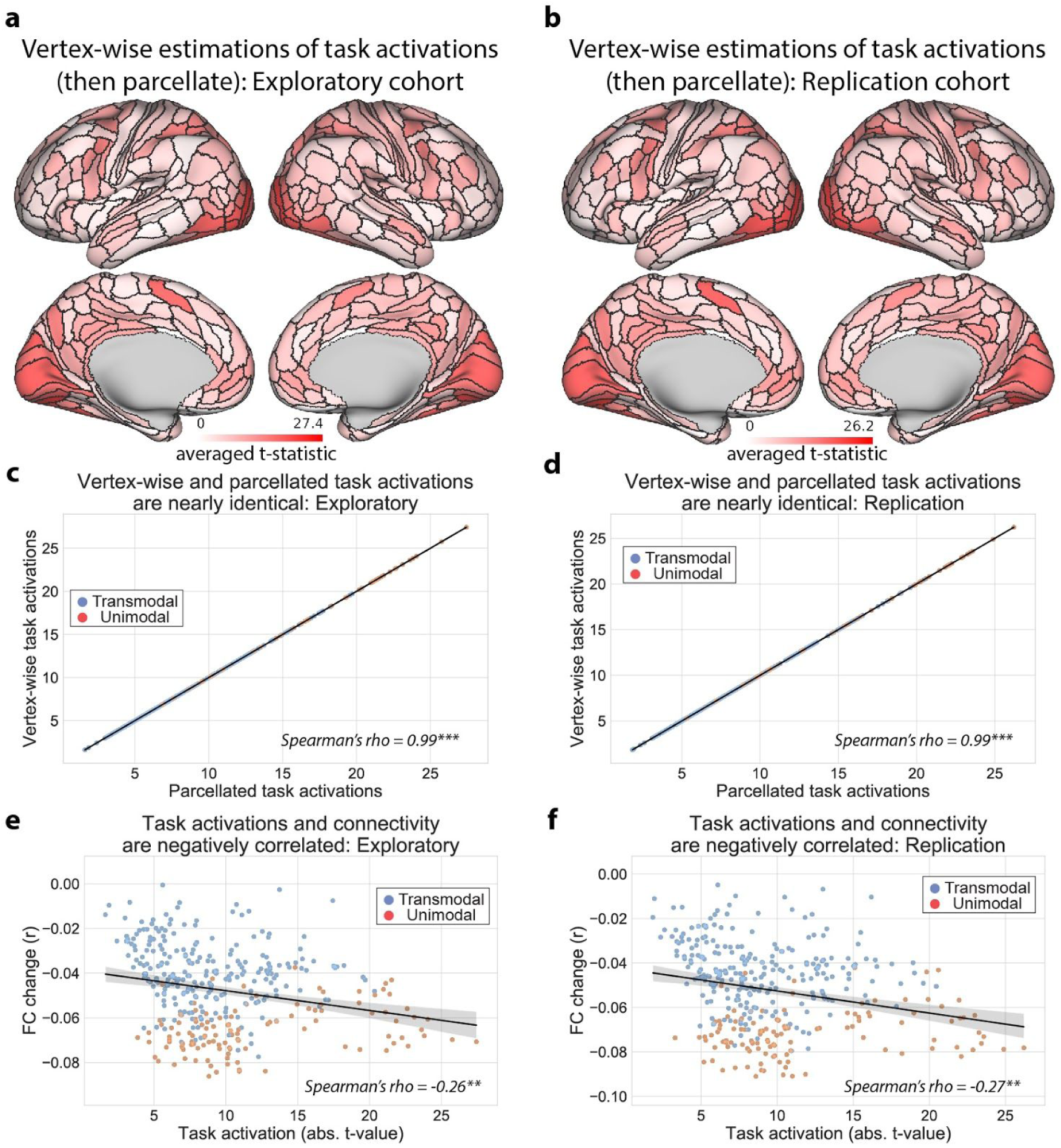
Comparison of using vertex-wise task GLM estimates versus parcellated task GLM estimates for task activation estimation. **a,b)** We estimated the task activation maps by first estimating the task GLM coefficients for each vertex, and then parcellating the data (i.e., estimate-then-parcellate) for both the exploratory and replication cohorts. **c,d)** We compared the task activation maps using the estimate-then-parcellate approach versus the parcellate-then-estimate (i.e., parcellate data and then perform task activation estimation). Overall, we found that the approaches yielded virtually identical task activation estimates. We note, however, that no nonlinear preprocessing steps (e.g., z-scoring of time series) were performed, thus making it unlikely that any differences would be observed. **e,f)** We performed the same analysis as in Figure 2, where we compared the task activation estimates using the estimate-then-parcellate approach with the FC change estimates. Again, we found virtually identical results. (*** = p<0.001, ** = p<0.01, * = p<0.05)

**Supplementary Figure 4.**
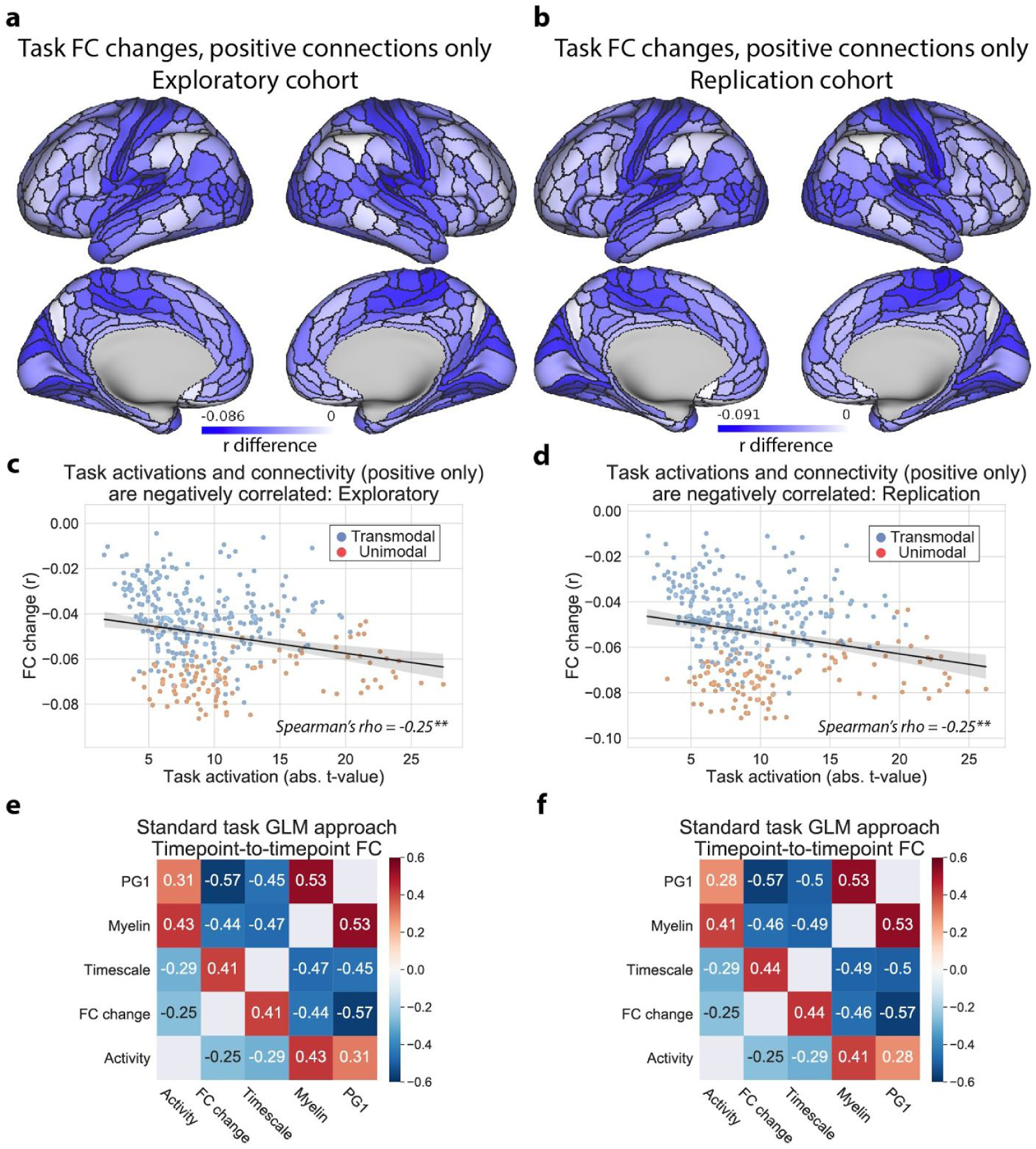
Comparison of results using thresholded (positive only) FC matrices. We computed FC change values (i.e., task versus rest) on thresholded resting-state and task-state FC matrices, removing any negative FC estimates. (This was applied on each rest- and task-state FC matrix independently.) We found virtually identical results, corroborating our findings that task-state FC reductions are most prominent in sensorimotor areas, and reductions follow a hierarchical gradient. **a,b)** The average task-state FC change for every brain parcel, for the exploratory and replication cohorts. **c,d)** The correlation across parcels between task-state FC change and task-evoked activation magnitudes. **e,f)** The summary correlation matrix, demonstrating that the associations between task-state FC changes and task activation, intrinsic timescale, myelin, and PG1 maps are virtually unchanged when using positive FC values only. (Note that correlations between variables, such as myelin and timescale, will be unchanged from Figure 5, since the only changed variable here was the FC change map.)

